# Classifying Calcium Imaging Dynamics with Deep Learning: Multi-Frequency Analysis through Quantile-Based Time-Series Network Representations

**DOI:** 10.64898/2025.12.08.693018

**Authors:** Caroline L. Alves, Simone Hufgard, Margot Mayer, Helena Dasch, Andriana S. L. O. Campanharo, Loriz Francisco Sallum, Francisco A. Rodrigues, Christiane Thielemann

## Abstract

To address the limitations of calcium imaging data, we propose a segmentation-agnostic deep learning framework that integrates Quantile-Based Time-Series Network (QTN) representations with convolutional neural networks to classify neuronal dynamics across multiple spatial resolutions and acquisition frequencies. By transforming fluorescence traces into compact, fixed-size matrices derived from quantile transitions, the method standardizes inputs across recordings while markedly reducing dimensionality and computational cost. Several QTN variants were systematically evaluated, demonstrating strong and consistent classification performance across both whole-image and grid-based preprocessing strategies. Notably, the framework maintained high accuracy under reduced temporal resolution and controlled noise perturbations, confirming that discrimination arises from meaningful temporal patterns rather than artifacts. This study establishes a robust, scalable, and generalizable approach for analyzing calcium imaging dynamics, paving the way for efficient, segmentation-independent characterization of neuronal activity in pharmacological and systems neuroscience applications.

## I. INTRODUCTION

### A. Challenges in Calcium Imaging Analysis

Calcium imaging provides a powerful, non-invasive technique for monitoring neuronal population activity at high spatial and temporal resolution [1, 2]. The recording of intracellular calcium transients enables the observation of network functionality and the impact of neuromodulatory factors on neuronal communication. However, calcium imaging data are inherently complex to analyze due to the indirect nature of calcium signals as proxies for action potentials, their temporal delay, and the presence of motion artifacts, shot noise, and fluorescence overlap in densely packed cellular environments [3–5].

These challenges complicate segmentation, signal extraction, and the subsequent classification of neuronal activity patterns—especially in low-contrast or noisy recordings.

Various methods have been developed to process calcium imaging data, ranging from classical signal-processing and statistical approaches [6, 7] to modern deep learning frameworks [8]. Convolutional neural networks (CNNs), in particular, have revolutionized biological image analysis by learning hierarchical features directly from raw microscopy data [9–13]. They have achieved state-of-the-art results in diverse microscopy applications, including phenotype prediction from unlabeled bright-field images [14] and high-content screening [15]. In calcium imaging, supervised CNNs can reach near-human accuracy in neuron identification, outperforming manual annotation and unsupervised approaches such as PCA/ICA [16, 17]. By improving detection accuracy and reducing manual effort, CNNs have significantly accelerated large-scale calcium imaging analysis pipelines.

Nevertheless, conventional CNN-based methods still face major limitations when applied directly to calcium imaging. Models trained on static frames tend to prioritize spatial features while underutilizing the temporal structure that is essential to characterize neuronal dynamics [18, 19]. Furthermore, the segmentation step—often required to isolate single-cell activity—can introduce substantial variability and bias, especially in dense or low-signal datasets. These difficulties hinder the development of robust, scalable workflows for comparative and quantitative analyses of neuronal activity.

### B. Quantile-Based Time-Series Networks representations

A promising direction for capturing temporal structure without relying on explicit segmentation arises from the framework of time-series-to-network mappings [20–22].

For example, in [23], the concept of a duality between time series and networks was introduced, demonstrating that dynamical patterns can be represented as networks whose nodes correspond to discrete states of the signal and whose edges encode transitions between successive states. This representation preserves essential dynamical information—such as stationarity, correlation structure, and complexity—while transforming irregular or noisy time series into analyzable topologies.

A key step in this framework is determining the number of quantile partitions *Q*, which can be empirically estimated using the following scaling heuristic defined in [24]:

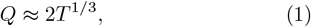

where *T* denotes the signal length. This empirical rule provides an effective balance between temporal resolution and statistical robustness. Furthermore, this work extended the duality concept through the *quantile graph approach* [24], where the continuous amplitude domain of a signal is partitioned into quantile bins, and transitions between these bins form a directed and weighted network.

Because quantile discretization is robust to noise and amplitude scaling, the resulting transition matrices capture relative variations and temporal dependencies in a compact and computationally efficient form. Building upon this principle, subsequent studies have developed quantile-based network models for dimensionality reduction and multivariate time-series analysis of diverse systems, including physiological, behavioral, and environmental datasets [25–29], demonstrating that network representations derived from quantile transitions can effectively encode temporal dynamics across complex datasets. However, this quantile-based network framework has not yet been applied to calcium imaging data, motivating their adoption in the present study.

### C. Aim and Innovation

Building on these developments, we propose a segmentation-agnostic framework that integrates Quantile-based Time-series Networks representations (QTNs) with CNN classification.

QTNs transform one-dimensional fluorescence traces into compact, fixed-size matrices by discretizing amplitudes into quantile bins and encoding transition probabilities between consecutive bins.

This transformation yields an interpretable, image-like representation of neuronal dynamics that preserves both temporal ordering and transition structure while substantially reducing data dimensionality and computational cost. By design, the approach provides a consistent input format across recordings with different acquisition rates, spatial resolutions, or signal-extraction strategies, thereby bypassing the need for explicit cell segmentation and its associated variability.

As a proof of concept, we apply this framework to neuronal networks exposed to brain-derived neurotrophic factor (BDNF)—a neurotrophin known to modulate synaptic plasticity, learning, and memory processes [30– 32].

Because BDNF has been extensively studied for its impact on neuronal activity and network reorganization [33], it serves as an ideal test case for validating our framework’s sensitivity to subtle, biologically relevant changes in network dynamics.

To our knowledge, this is the first study to employ QTN representations for calcium imaging data and the first to combine QTNs with CNNs to classify neuronal activity under neuromodulatory influence.

The proposed framework provides a generalizable, scalable, and segmentation-independent strategy for analyzing complex neuronal dynamics, offering a new methodological direction for high-dimensional calcium imaging datasets where conventional segmentation approaches remain unreliable.

## II. DATA COLLECTION

Primary rat cortical neurons (R-CX-500, Lonza, Switzerland) were cultured according to the QBM Cell Science Rat Cortical Neurons protocol provided by Axion Biosystems, with minor modifications. Briefly, cryopreserved cells were thawed at 37°C and gently resuspended in Neurobasal™ Plus Medium (Thermo Fisher Scientific) supplemented with 2% Neurobasal™ Plus Supplement, 0.5 mM GlutaMAX, and 1% penicillin/streptomycin (all from Thermo Fisher Scientific). Cells were plated on polyethyleneimine (PEI, Sigma-Aldrich, St. Louis, USA) and laminin (Sigma-Aldrich) coated ibidi dishes (Ibidi, Gräfelfing, Germany) at a density of 80000 cells in a 5 µL droplet and maintained at 37°C in a humidified 5% CO_2_ atmosphere. Half of the medium was replaced three times per week. As a positive control for the workflow, cultures were treated with brain-derived neurotrophic factor (BDNF; 50 ng/mL) at day in vitro (DIV) 8 for 72 hours.

For calcium imaging, samples were incubated with 5 µM Calbryte™ 520 AM (AAT Bioquest, Inc.) dissolved in NB+ medium at 37°C for 60 minutes. After incubation, the dye solution was replaced with fresh medium to remove excess probes. Before imaging, samples were maintained in the incubation chamber of the microscope at 37°C for an additional 15 minutes.

Calcium imaging was performed on a Leica DMI8-CS inverted confocal microscope (Stellaris 5) using an HC PL APO 10×/0.40 CS2 objective. Excitation wavelength was set to 493 nm, and laser power was minimized to reduce phototoxicity. Video sequences were acquired at 4 Hz (analog mode) or 28 Hz (fast photon-counting mode) with a resolution of 512 × 512 pixels and and a pixel ratio close to 1.

Calcium imaging recordings were performed three times per week, starting seven days after BDNF treatment. Each experimental group included a total of eight independent samples. For each culture, up to 12 individual fields of view (FOVs) were randomly selected across the network, resulting in 96 videos per condition for both the BDNF-treated and control groups.

To assess reproducibility and generalization, a second independent experiment was conducted using new BDNF-treated and control samples prepared with the same protocol and imaging parameters as the first. The recordings for this experiment started one day after the BDNF treatment. During the first week, there were daily imaging sessions, and after that, there were three sessions per week. This design yielded a total of 128 videos per condition. The recordings, acquired from non-overlapping FOVs, formed a separate dataset used exclusively for external validation, independent of model training or tuning.

## III. DATA PREPROCESSING

All calcium imaging recordings were preprocessed using a standardized pipeline to ensure consistent and reliable extraction of neuronal activity at multiple spatial scales, whose two complementary strategies to extract time series from the video data are: (i) Whole-Image Averaging (Section III A), which computes a global intensity trace by averaging pixel values across the entire field of view (FOV), capturing the overall population activity without spatial resolution; and (ii) Grid-Based Partitioning (Section III B), which divides the imaging field into non-overlapping regions to obtain spatially localized signals while avoiding the need for explicit cell segmentation.

For the grid-based approach, a two-step procedure was applied to select regions that exhibit meaningful neuronal activity. First, the fluorescence time series of each spatial unit grid was evaluated for temporal variability using the interquartile range (IQR), a robust measure that highlights signal fluctuations while minimizing the influence of outliers. Next, we restricted the analysis to spatial regions whose mean pixel intensities fell within the 25th to 75th percentile of the global intensity distribution, corresponding to gray-level ranges where functional neuronal activity is most likely to occur. This percentile-based filtering method—commonly used in computer vision to isolate regions with significant intensity variation—excludes saturated, low-signal, or artifact-prone areas and prioritizes regions exhibiting biologically relevant dynamics [34].

### A. Whole-Image Averaging

In the whole-image averaging approach, a single fluorescence time series was generated by averaging the pixel intensities of each frame in the calcium imaging video.

Although this procedure removes spatial information—which is often the main advantage of calcium imaging—it was intentionally included here as a baseline representation to evaluate the performance and robustness of the proposed QTN framework under the most spatially agnostic condition.

This global trace captures the overall temporal evolution of network activity across the entire field of view (FOV), allowing direct assessment of whether QTN-based representations and CNN classification can still detect biologically meaningful differences when spatial heterogeneity is collapsed. By comparing the results obtained from the whole-image approach with those from the grid-based analysis (subsection III B), we quantified how spatial resolution influences the discriminative power of QTN-derived features.

### B. Grid-Based Partitioning

In the grid-based partitioning approach, each calcium imaging frame was divided into non-overlapping 16 × 16-pixel grids, enabling localized extraction of neuronal activity signals across the FOV. For each grid, multiple distinct fluorescence time series were extracted, each capturing a different aspect of the local dynamics. These included: (i) the average fluorescence intensity over time (AIP), representing the mean activity level within the region; (ii) the first (DAIP) and second temporal derivatives (DAIP), which describe the rate and acceleration of fluorescence changes; (iii) the coefficient of variation (CV), a widely used indicator of relative signal variability normalized by mean intensity [35, 36]; (iv) signal entropy, which quantifies the irregularity and complexity of fluorescence fluctuations [37, 38]; and (v) the low-frequency spectral power. The low-frequency power (*P*_LF_) was computed from the real-valued fluorescence time series using the real fast Fourier transform (rFFT), defined as

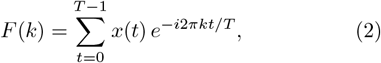

where *x*(*t*) is the fluorescence intensity at time *t, T* is the total number of frames, and *F* (*k*) denotes the complex Fourier coefficient corresponding to the *k*-th frequency bin. The low-frequency spectral power was then calculated as

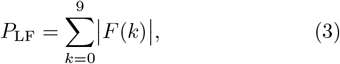

which represents the sum of the magnitudes of the first ten frequency components, including the DC term. The frequency resolution was defined as Δ*f* = *f*_*s*_*/T*, where *f*_*s*_ is the sampling rate and *T* is the number of frames in the recording. This corresponds to Δ*f ≈* 0.0087 Hz for analog recordings acquired at *f*_*s*_ = 4 Hz with *T* = 460 frames, and Δ*f ≈* 0.0083 Hz for fast recordings acquired at *f*_*s*_ = 28 Hz with *T* = 3356 frames. Consequently, bins *k* = 0, …, 9 capture oscillations below approximately 0.08 Hz, summarizing slow oscillatory components associated with coordinated neuronal activity [39, 40].

Although the same features could in principle be computed for the whole-image signal described in Section III A, we restricted them to the grid-based approach because local dynamics are otherwise averaged out in global representations.

Each of these time series was subsequently converted into a QTN (described in the subsection IV A), producing a compact matrix that preserves temporal ordering and transition dynamics. The resulting QTN matrices—one per descriptor and per grid—were then used as inputs to the CNN classifier. This multi-descriptor approach provides a spatially resolved and physiologically meaningful representation of neuronal activity, enhancing sensitivity to functional differences while remaining independent of explicit cell segmentation.

## IV. METHODOLOGY

### A. QTN Representations

To extract robust descriptors of temporal dynamics from calcium imaging data, we used Quantile-based Transition Network (QTN) representations, which convert each fluorescence time series into a structured, image-like matrix. QTNs operate on individual time series and can therefore be applied uniformly across all signal extraction strategies used in this study—namely, Whole-Image Averaging (Section III A) and Grid-Based Partitioning (Section III B).

Four complementary representations were generated for each time series: Symbolic Aggregate approXimation (SAX) Transition Graphs, Quantile Graphs (QG), Gramian Angular Fields (GAF), and Markov Transition Fields (MTF). The number of quantile bins (*Q*) was determined using Eq. 1, which balances temporal resolution and statistical robustness while maintaining the Hurst exponent stable across signals of different lengths. Accordingly, *Q* was set to 15 for recordings acquired at 4 Hz (460 frames) and 30 for recordings acquired at 28 Hz (3356 frames).

- **SAX Transition Graphs [41]:** The Symbolic Aggregate approXimation (SAX) method discretizes each time series into *Q* equiprobable bins after z-score normalization, converting the signal into a sequence of symbols that capture its ordinal patterns. A directed transition graph is then generated by counting the frequency of symbol-to-symbol transitions between consecutive time points, resulting in a *Q* × *Q* adjacency matrix that compactly encodes the temporal dynamics of the signal while remaining robust to amplitude scaling and noise.
- **Quantile Graphs (QG)** [**26**, **27**]: Extending the SAX framework, QG matrices extend the analysis to multiple temporal lags (*k* = 1 to *k* = 20), capturing long-range dependencies and recurrent temporal patterns. This approach generates a series of adjacency matrices that encode transitions between quantile-defined states over varying time scales, providing a richer representation of the underlying dynamics. In our context, the QG method transforms fluorescence intensity time series into directed, weighted graphs whose topology captures transitions between quantile-defined amplitude states, as conceptually illustrated in Figure 1.
- **Gramian Angular Fields (GAF) [42]:** The GAF represents temporal correlations by mapping the normalized time series into a polar coordinate system and computing the cosine of the sum of pair-wise angles. This produces a symmetric *Q × Q* matrix that encodes smooth trends and cyclic patterns, providing a geometric representation of amplitude relationships over time.
- **Markov Transition Fields (MTF) [43]:** The MTF encodes temporal dynamics by first constructing a first-order Markov chain that captures transition probabilities between quantile bins [23]. These probabilities are then expanded into a *Q × Q* matrix, where each element reflects the likelihood of transitions between states at different time points, emphasizing the evolving dynamics of the signal throughout the recording.

Figure 2 provides examples of the four QTN representations derived from a representative region of interest (ROI) in a 4 Hz recording. All QTN matrices were standardized to uniform dimensions appropriate for subsequent CNN-based classification.

**FIG. 1:**
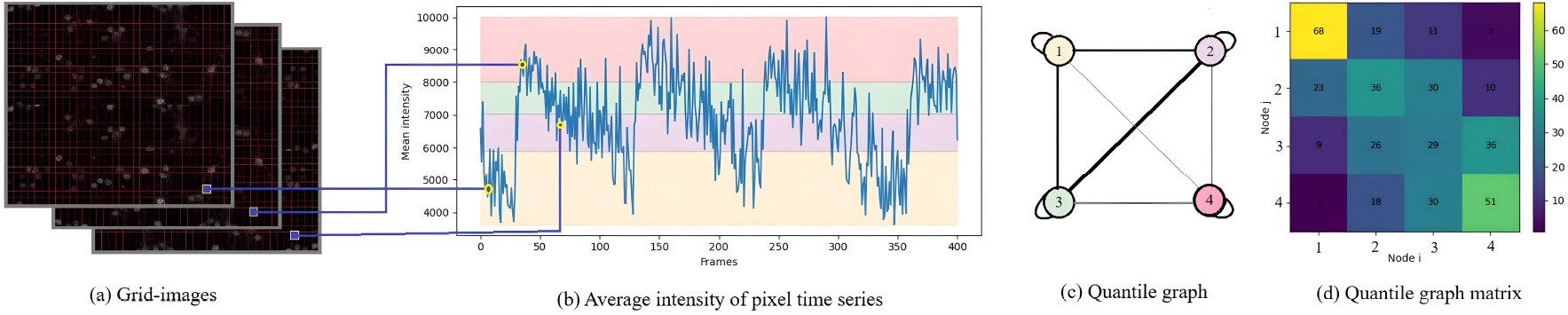
Conceptual illustration of the Quantile Graph (QG) construction process. (a) Calcium imaging frames are partitioned into spatial grids, from which fluorescence intensity time series are extracted. (b) A representative trace displays the average pixel intensity across frames, subdivided into four quantile ranges for clarity in this explanation. (c) Each quantile interval defines a node, and transitions between successive quantiles (k = 1) establish directed, weighted edges that collectively form the Quantile Graph. (d) The corresponding adjacency matrix summarizes these transition frequencies. This scheme illustrates how temporal dynamics are transformed into a compact network representation, with each node corresponding to a quantile-defined state and each edge representing a transition between them. Abbreviations: QG, Quantile Graph.

**FIG. 2:**
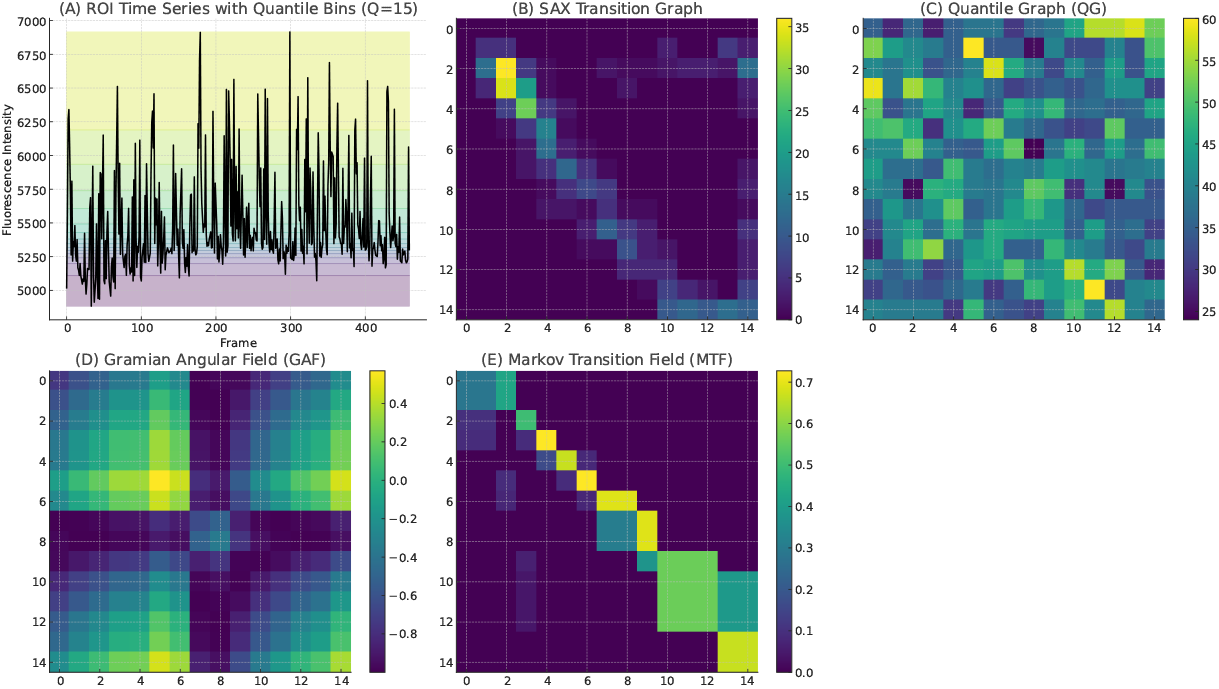
Illustration of QTN-based representations. (A) Example of a randomly selected region of interest (ROI) extracted from a calcium imaging recording acquired at 4 Hz (460 frames). The fluorescence intensity time series is shown with quantile bins (*Q* = 15) highlighted in distinct colors. From this time series, four QTN matrices of uniform size 15 × 15 were computed: (B) SAX, (C) QG, (D) GAF, and (E) MTF. Abbreviations: ROI, Region of Interest; QTN, Quantile-based Transition Network; SAX, Symbolic Aggregate approXimation; QG, Quantile Graph; GAF, Gramian Angular Field; MTF, Markov Transition Field.

In the grid-based approach, each imaging frame was divided into non-overlapping 16×16-pixel regions, and a fluorescence time series was extracted from each grid by averaging the pixel intensities within that region. QTN matrices were then computed for every grid to capture the temporal dynamics of localized activity (Figure 3).

**FIG. 3:**
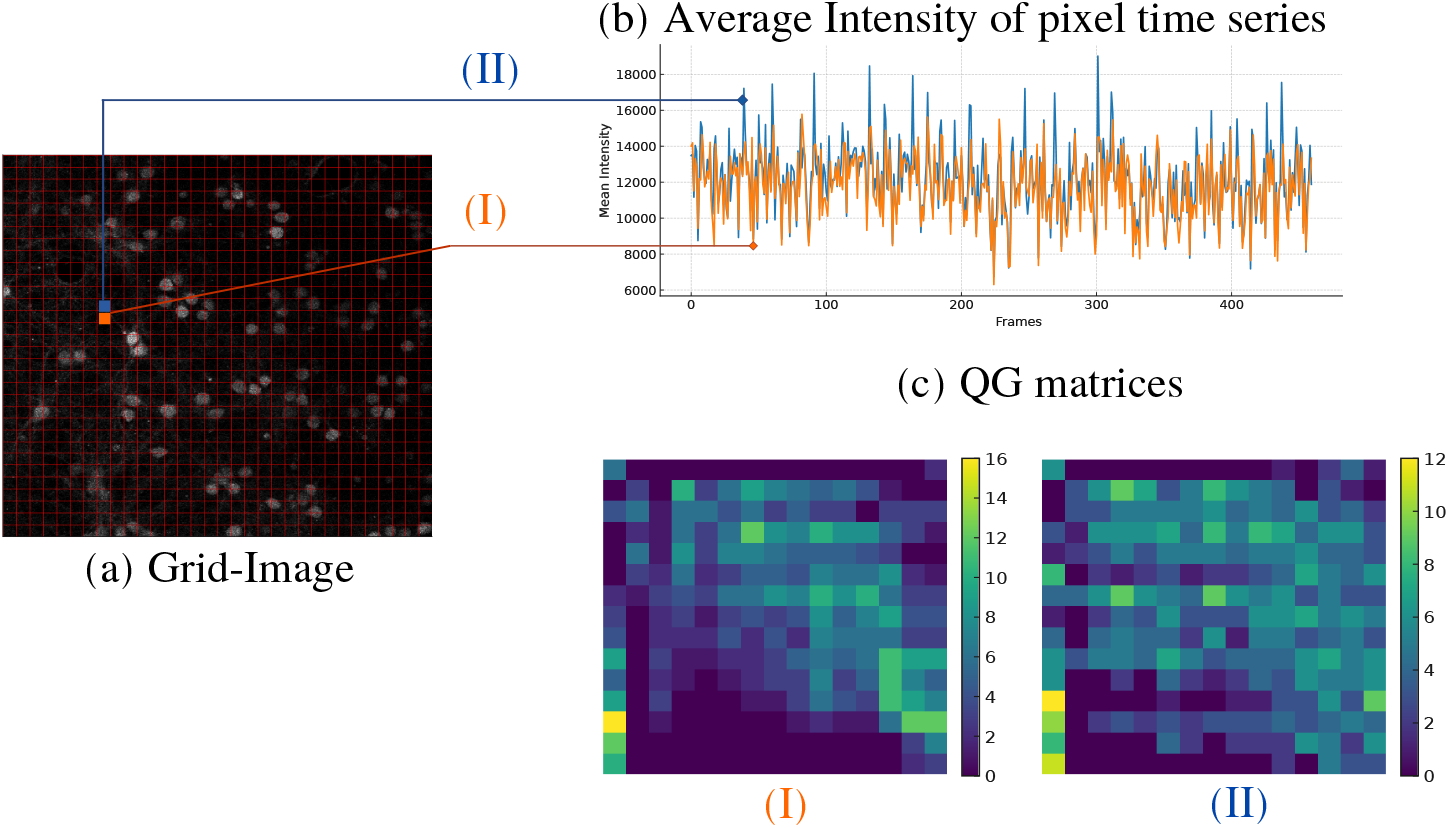
Workflow for QTN representations from grid-partitioned calcium imaging. (a) The original calcium imaging frame is divided into a 16×16 grid of spatial units. Two representative regions, highlighted in orange (I) and blue (II), are shown for illustration. (b) For each selected grid, the mean fluorescence intensity is computed across frames, producing localized time series that reflect neuronal activity. (c) Corresponding QG matrices derived from time series (I) and (II), with matrix colors matching the original grid selections in (a) and (b). Abbreviations: QG, Quantile Graph; QTN, Quantile-based Transition Network.

### B. CNN

To classify BDNF-treated samples against SHAM controls, we trained a Convolutional Neural Network (CNN) using structured matrices derived exclusively from Quantile-based Time-series (QTN) representations of calcium imaging data.

Each extracted fluorescence time series—obtained through Whole-Image Averaging or Grid-Based Partitioning—was converted into a two-dimensional QTN matrix that encodes the temporal transition dynamics of the signal.

These matrices served directly as image-like inputs to the CNN, enabling the model to learn discriminative spatiotemporal patterns in neuronal activity without requiring handcrafted features or segmentation-based connectivity estimation.

As described in Subsection IV A, for each acquisition frequency, all QTN matrices were standardized to the same fixed size, ensuring uniform input dimensions for subsequent CNN-based classification. These matrices served directly as image-like inputs to the CNN, enabling the model to learn discriminative spatiotemporal patterns in neuronal activity without requiring handcrafted features or segmentation-based connectivity estimation. Hyperparameter optimization was conducted using random search, implemented via KerasTuner [44–46], to systematically explore a predefined search space. The CNN architecture and hyperparameter search space are detailed in Table I. To manage computational resources, the optimization process was limited to 1 trial with 2 executions per trial, and preliminary exploration was conducted over 20 epochs to identify promising configurations. Final model training was executed for up to 2000 epochs.

To enhance training efficiency and model generalization, early stopping was applied to halt training if the validation loss failed to improve over 100 consecutive epochs [47, 48]. Additionally, a learning rate scheduler was employed to dynamically reduce the learning rate upon detection of validation loss plateaus [49], thereby mitigating computational overhead while allowing fine-grained adjustments during later training phases.

For model training and evaluation, the QTN matrices generated from all recordings served as the inputs to the CNN. The dataset matrices were initially partitioned into training (75%) and testing (25%) subsets using a stratified holdout split, which preserved the original class distribution of BDNF-treated and SHAM samples to ensure balanced representation and mitigate sampling bias. Within the training set, a further 30% validation split was employed during training to monitor generalization performance. No samples or recordings from the second experiment were used during model training, ensuring complete independence between the training and validation datasets.

To visualize model performance, Receiver Operating Characteristic (ROC) curves were employed, with the area under the curve (AUC-ROC) serving as a key metric, ranging from 0.5 (random classification) to 1 (perfect discrimination) [50, 51]. Accuracy was used as the primary evaluation metric, quantifying the overall correctness of predictions across all classes [52–55]. Additionally, precision and recall were computed to assess class-specific performance [56–58]; precision measures the proportion of correctly predicted SHAM samples among all SHAM predictions, while recall (sensitivity) evaluates the model’s ability to correctly identify BDNF-treated samples. The random rearch hyperparameter optimization process was guided by validation accuracy, ensuring that the selected models prioritized overall classification correctness.

## V. RESULTS

### A. Whole-Image Averaging

To assess the impact of acquisition frequency on classification performance, we compared 4 Hz and 28 Hz frequency recordings using QTN representations as input to CNN models. As shown in Figure 4, 28 Hz frequency recordings consistently achieved superior AUC values across all QTN methods, particularly enhancing the discriminative power of SAX and MTF representations. Nonetheless, 4 Hz frequency recordings also yielded strong performance, maintaining high AUCs, which demonstrates their effectiveness despite the lower temporal resolution. Figure 5 further illustrates this trend through a heatmap of classification accuracies comparing 4 Hz and 28 Hz acquisition frequencies. Furthermore, to further assess classification discriminability, ROC curves were generated for both 4 Hz and 28 Hz acquisition frequency recordings across all QTN methods. As depicted in Figure 6, for 28 Hz acquisition frequency recordings, all methods exhibited near-perfect ROC profiles, with AUC values approaching 1.00, including SAX (AUC = 0.98), QG (AUC = 0.99), GAF (AUC = 1.00), and MTF (AUC = 0.99). For 4 Hz acquisition frequency recordings, although slightly lower, also demonstrated a strong classification capacity, with AUC values of 0.94 (SAX), 0.97 (QG), 1.00 (GAF), and 0.88 (MTF), underscoring their robustness despite their lower temporal resolution.

**FIG. 4:**
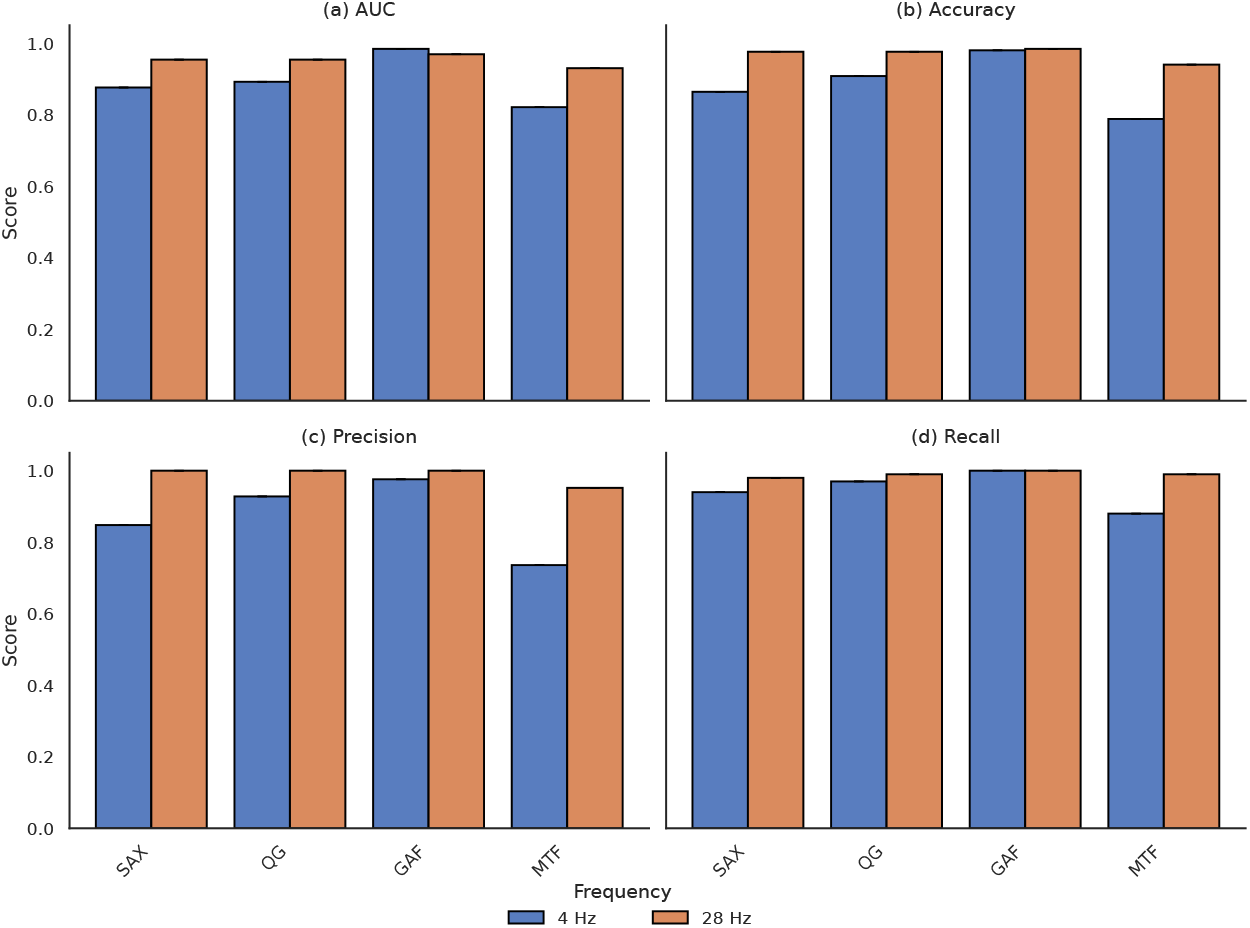
Whole-image averaging comparison of classification performance across acquisition frequencies. Comparison of classification performance between 4 Hz and 28 Hz frequency recordings using QTN Representations. The plots display (a) AUC, (b) Accuracy, (c) Precision, and (d) Recall across QTN methods (SAX, QG, GAF, MTF). AUC: Area Under the Curve; QTN: Quantile-based Transition Network; SAX: Symbolic Aggregate approXimation; QG: Quantile Graph; GAF: Gramian Angular Field; MTF: Markov Transition Field.

**FIG. 5:**
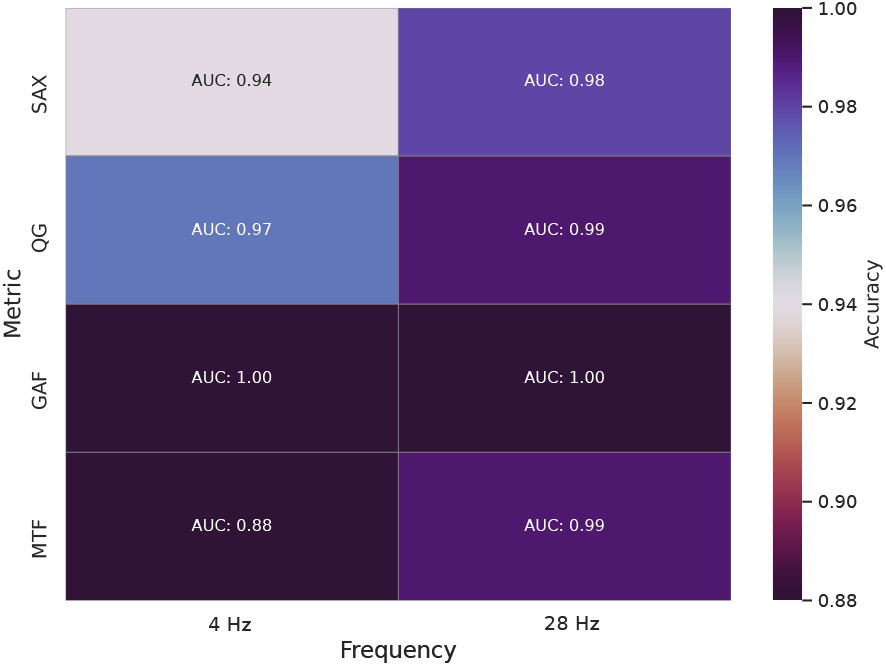
Whole-image averaging heatmap of classification accuracies for 4 Hz and 28 Hz frequency recordings across QTN methods. Annotated values represent corresponding AUCs. QTN: Quantile-based Transition Network; AUC: Area Under the Curve.

**FIG. 6:**
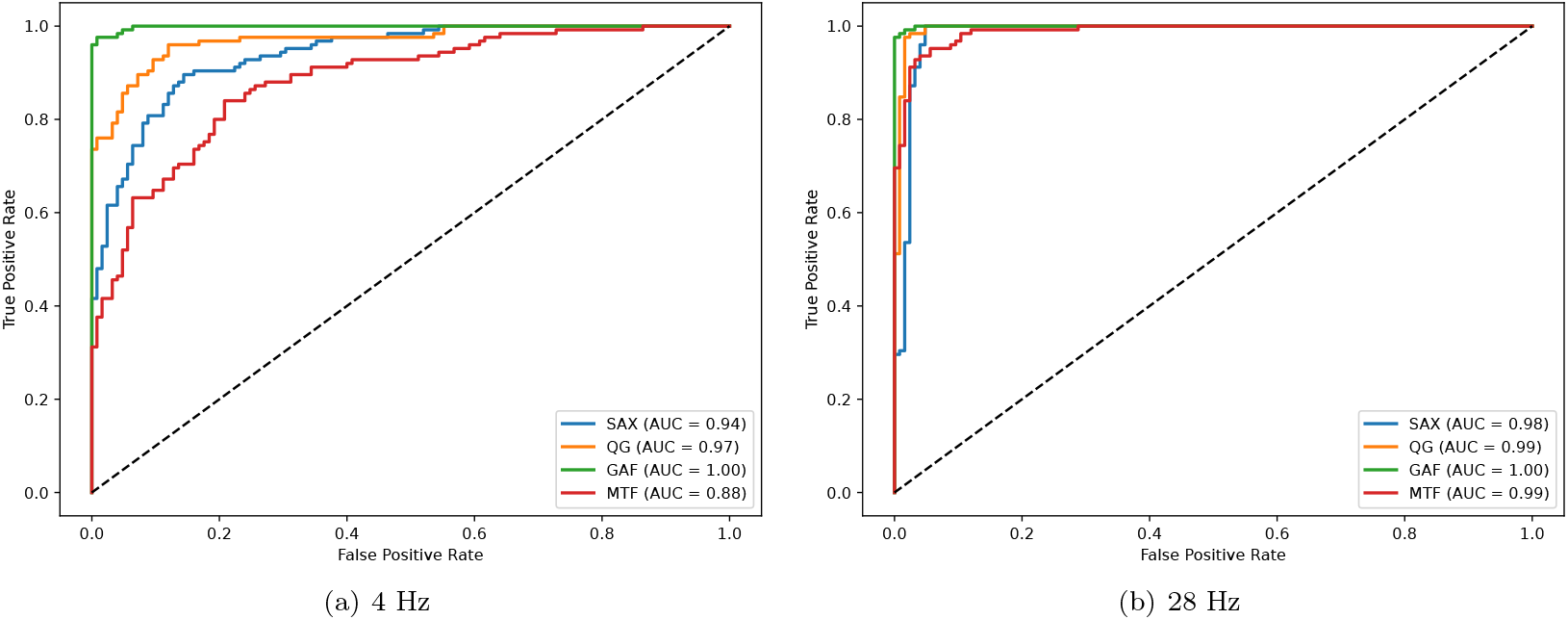
Whole-image averaging ROC curves for 4 Hz and 28 Hz frequency recordings across QTN methods. ROC curves comparing the classification performance of QTN metrics for 4 Hz and 28 Hz frequency recordings. Annotated AUC values indicate superior classification performance for a 4 Hz acquisition frequency compared to 28 Hz, while still demonstrating robust performance at lower frequencies. QTN: Quantile-based Transition Network; ROC: Receiver Operating Characteristic; AUC: Area Under the Curve

The confusion matrices presented in Figure 7 provide a detailed breakdown of classification outcomes for each QTN method. For example, the SAX representation achieved high classification accuracy, correctly identifying 97% of BDNF-treated samples and 93% of SHAM controls, underscoring its robustness when applied to recordings at a 28 Hz acquisition frequency. In comparison, GAF maintained perfect classification performance across both conditions. By contrast, QG and MTF exhibited slightly higher off-diagonal values, indicating minor misclassifications yet preserving strong overall accuracy.

**FIG. 7:**
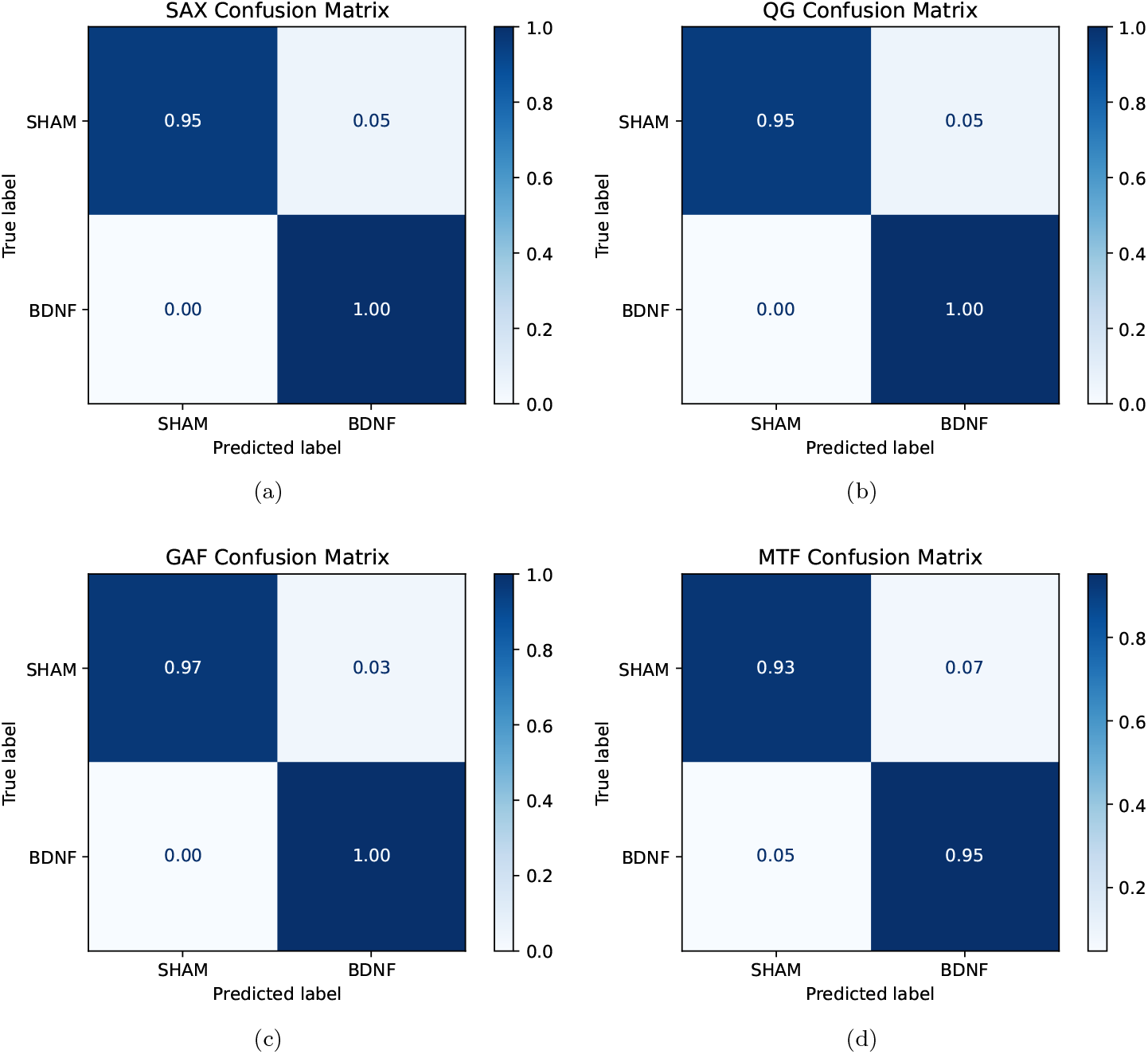
Whole-image confusion matrices for each QTN method (SAX, QG, GAF, MTF). The diagonal values indicate the proportion of correctly classified SHAM and BDNF samples. GAF achieved perfect classification for BDNF (1.00), with minimal SHAM misclassification (0.03). Values are normalized. QTN: Quantile-based Transition Network; SAX: Symbolic Aggregate approXimation; QG: Quantile Graph; GAF: Gramian Angular Field; MTF: Markov Transition Field; BDNF: Brain-Derived Neurotrophic Factor; SHAM: Control condition without treatment

The learning curves depicted in Figure 8 illustrate the training and validation accuracy trajectories across epochs for each QTN method. All methods demonstrated rapid convergence within the initial epochs, stabilizing with minimal overfitting as training progressed. Specifically, GAF (green) maintained consistently high validation accuracy throughout the training process, reflecting its robustness across iterations. SAX (blue) and QG (orange) exhibited stable learning dynamics, reaching high accuracy levels with minimal fluctuations. MTF (red) required slightly more epochs to stabilize but ultimately achieved comparable performance. The shaded regions surrounding each curve represent the standard deviation, capturing variability across training runs and indicating reliable generalization performance across all QTN representations.

**FIG. 8:**
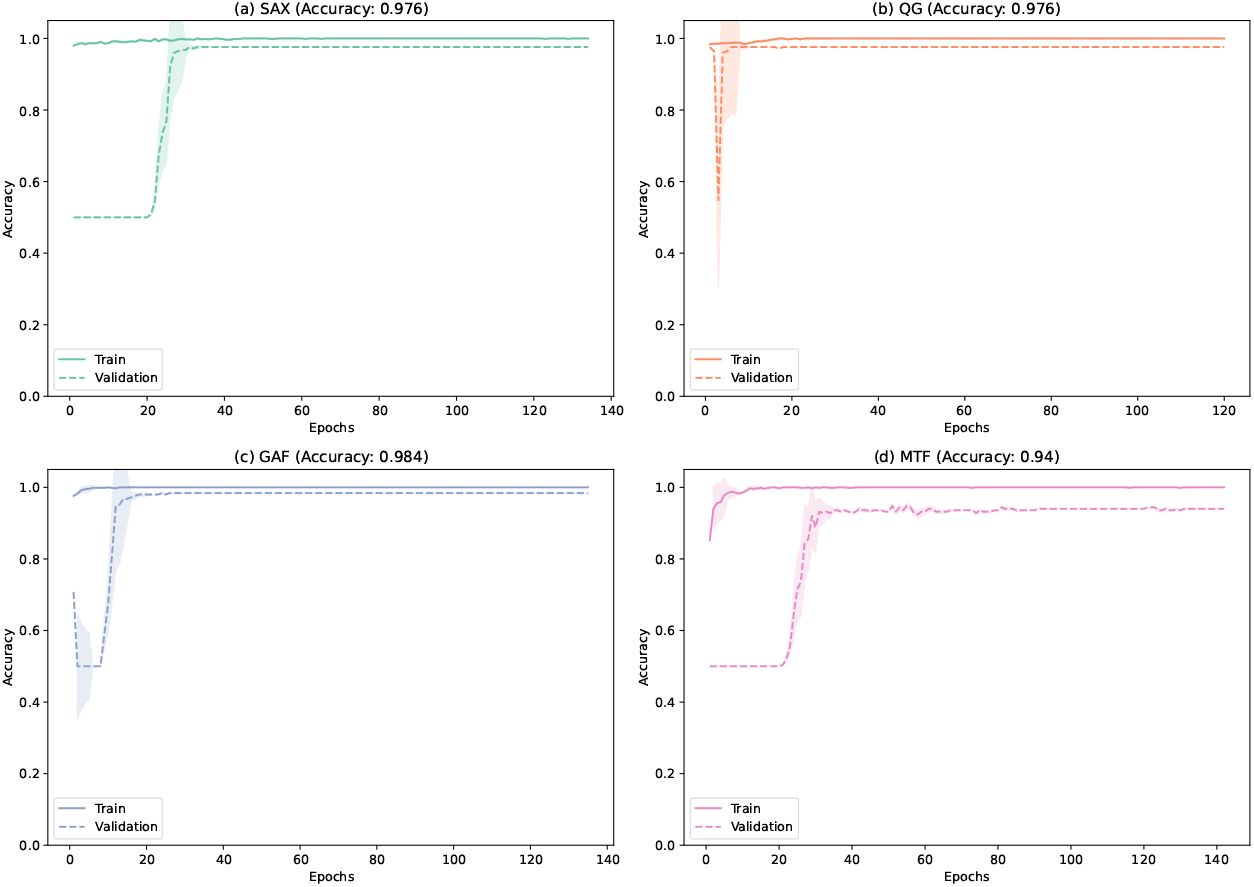
Whole-image averaging learning curves with shaded validation error across QTN methods for 28 Hz acquisition frequency recordings. Training (solid lines) and validation (dashed lines) accuracies are plotted across epochs for each QTN method: (a) SAX (blue), (b) QG (orange), (c) GAF (green), and (d) MTF (red). The shaded regions around each curve represent the standard deviation, indicating variability across training and validation iterations. QTN: Quantile-based Transition Network; SAX: Symbolic Aggregate approXimation; QG: Quantile Graph; GAF: Gramian Angular Field; MTF: Markov Transition Field.

To verify that the CNN’s perfect classification performance with GAF representations was not driven by noise memorization, we performed a robustness analysis by progressively adding Gaussian noise to the input matrices. This controlled perturbation assessed whether the model’s discriminative capacity relied on meaningful temporal patterns rather than artifacts. As shown in Supplementary VII in Figures S1–(a), the model sustained high AUC values (above 0.9) up to moderate noise levels (standard deviation *≈* 2.0), with performance gradually declining under heavier perturbations. These findings confirm that the CNN model learned robust signal structures, rather than overfitting to noise.

**TABLE I:**
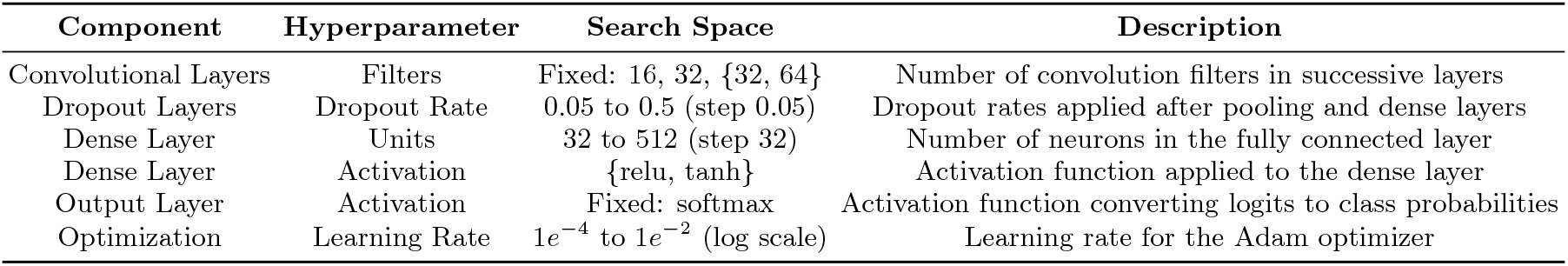
Hyperparameter search space explored for the CNN architecture during random search optimization. CNN: Convolutional Neural Network; relu: Rectified Linear Unit function; tanh: hyperbolic tangent function.

| Component | Hyperparameter | Search Space | Description |
| --- | --- | --- | --- |
| Convolutional Layers | Filters | Fixed: 16, 32, {32, 64} | Number of convolution filters in successive layers |
| Dropout Layers | Dropout Rate | 0.05 to 0.5 (step 0.05) | Dropout rates applied after pooling and dense layers |
| Dense Layer | Units | 32 to 512 (step 32) | Number of neurons in the fully connected layer |
| Dense Layer | Activation | {relu, tanh} | Activation function applied to the dense layer |
| Output Layer | Activation | Fixed: softmax | Activation function converting logits to class probabilities |
| Optimization | Learning Rate | $1e^{-4}$ to $1e^{-2}$ (log scale) | Learning rate for the Adam optimizer |

### B. Grid-Based Partitioning

Building on the grid-based segmentation approach described in Section III B, we evaluated how different feature descriptors influence classification performance when processed through QTN representations and fed into CNN models. The results, summarized in Figure 9, demonstrate that AIP and Shannon entropy (Entropy) features consistently yielded superior classification performance across all QTN methods, as reflected in higher AUC, accuracy, precision, and recall scores.

**FIG. 9:**
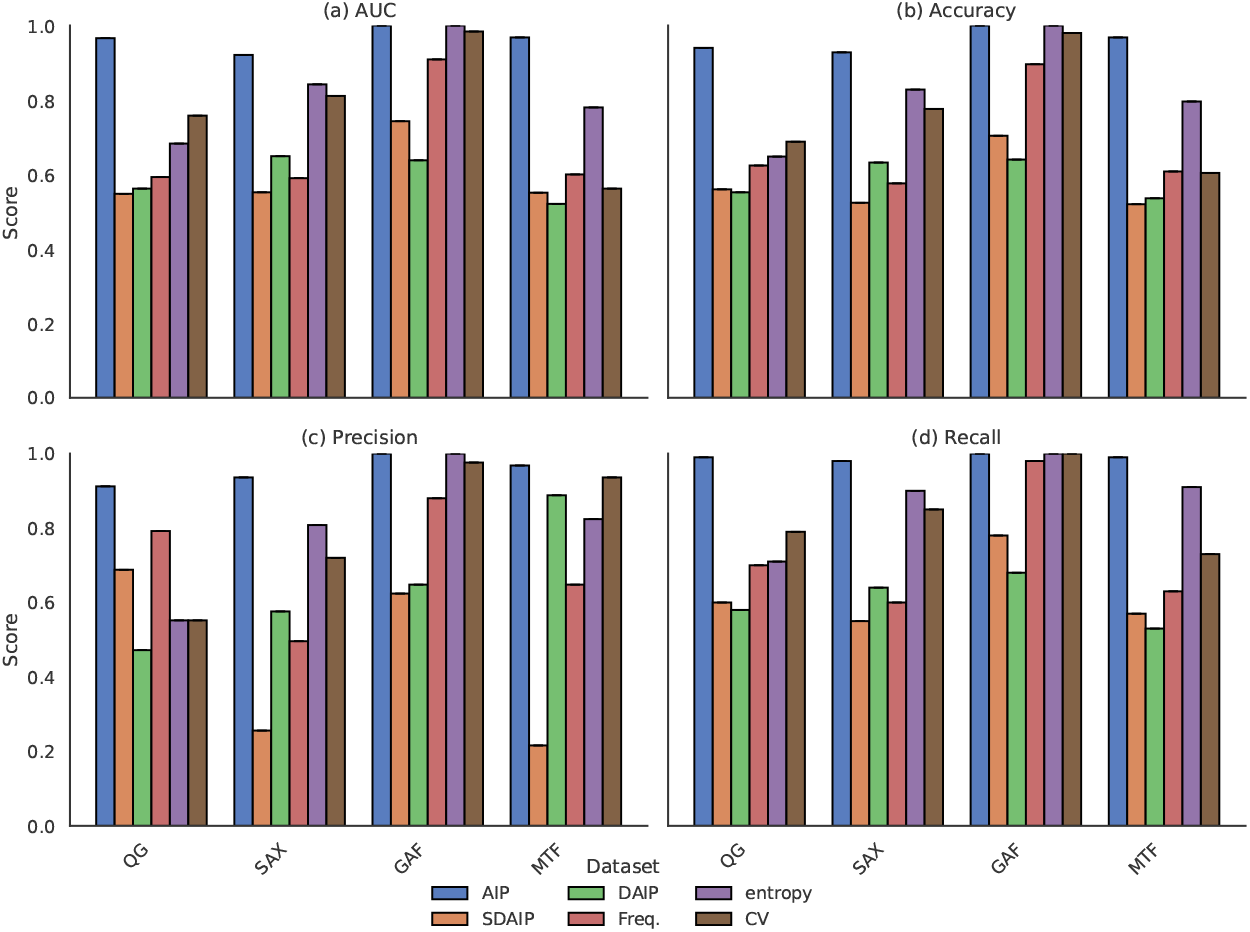
Grid-based partitioning comparison of classification performance across feature descriptors using 28Hz. Classification performance using QTN Representations applied to spatially localized grid-derived features: (a) AIP, (b) DAIP, (c) SDAIP, (d) Entropy (Shannon entropy), (e) Low-Frequency Spectral Power (Freq.), and (f) CV. The plots display (a) AUC, (b) Accuracy, (c) Precision, and (d) Recall across QTN methods (SAX, QG, GAF, MTF). QTN: Quantile-based Transition Network; AIP: Averaged Intensity Pixel; DAIP: Derivative of AIP; SDAIP: Second Derivative of AIP; Freq.: Low-Frequency Spectral Power; CV: Coefficient of Variation; SAX: Symbolic Aggregate approXimation; QG: Quantile Graph; GAF: Gramian Angular Field; MTF: Markov Transition Field; AUC: Area Under the Curve.

The strong performance of AIP suggests that absolute fluorescence intensity fluctuations retain critical information regarding global and local activity changes induced by BDNF treatment. Entropy (Shannon entropy), on the other hand, effectively captures the irregularity and complexity of intensity variations within each grid, providing a robust descriptor of dynamic signal heterogeneity. In contrast, derivative-based features—first (DAIP) and second derivatives (SDAIP)—emphasize rapid signal changes and are more susceptible to amplifying noise, resulting in lower classification performance. Features derived from low-frequency spectral power (Freq.) and coefficient of variation (CV) exhibited intermediate performance, balancing sensitivity to signal amplitude and temporal stability.

The classification outcomes for each QTN method using grid-partitioned AIP time series are detailed in Figure 10. Notably, the GAF representation achieved perfect classification, correctly identifying all BDNF-treated and SHAM samples without misclassifications. MTF also demonstrated high discriminative capacity, with balanced classification rates of 97% for both classes. SAX and QG exhibited slightly higher off-diagonal values but still maintained strong overall accuracy, confirming the effectiveness of spatially localized signal decomposition in enhancing classification performance.

**FIG. 10:**
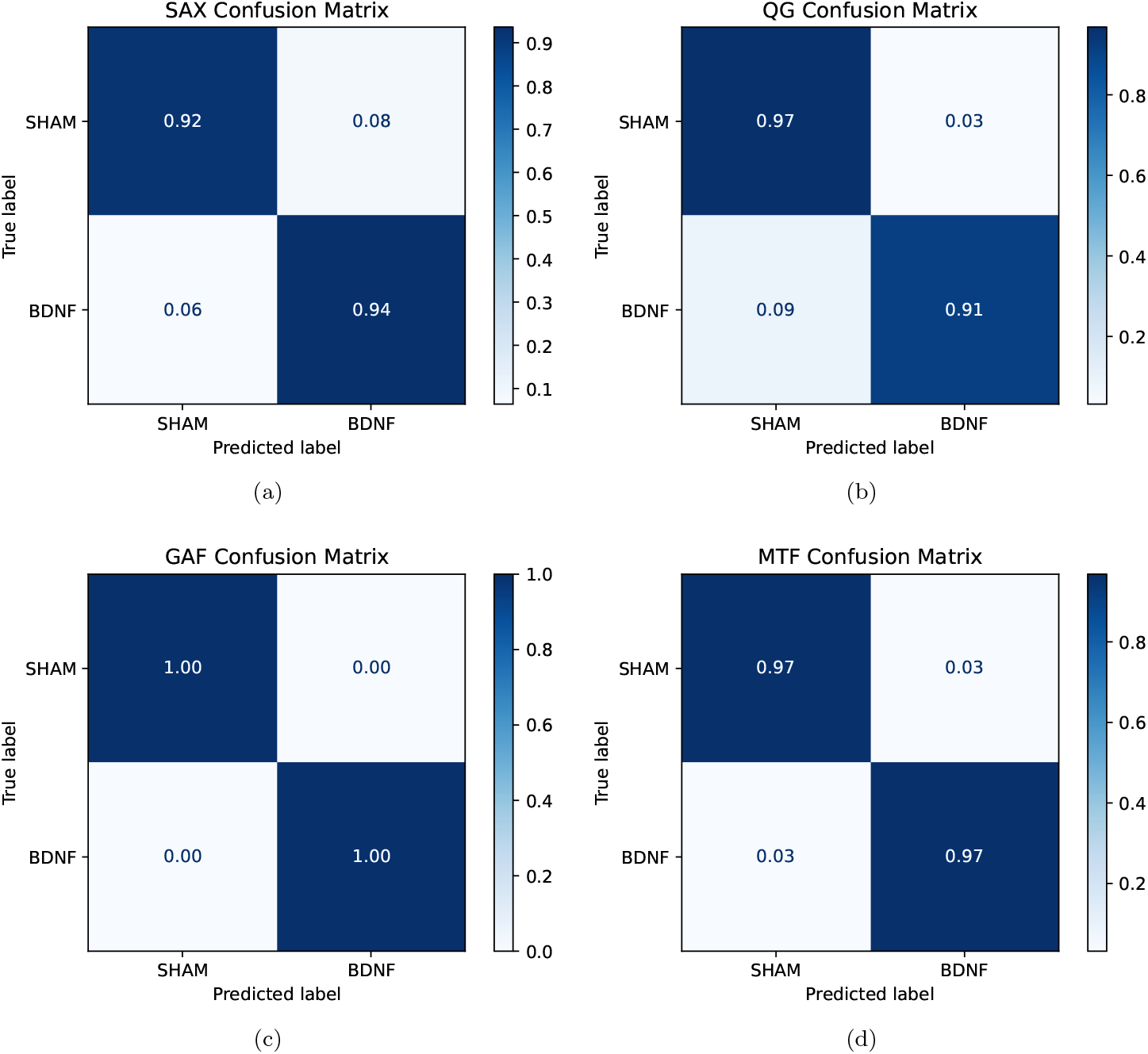
Grid-based partitioning confusion matrices for each QTN method applied to grid-partitioned AIP time series using 28 Hz. GAF achieved perfect classification for both SHAM and BDNF samples (1.00). SAX, QG and MTF also showed strong performance, with slight off-diagonal values indicating isolated errors. Values are normalized. QTN: Quantile-based Transition Network; AIP: Averaged Intensity Pixel; GAF: Gramian Angular Field; SAX: Symbolic Aggregate approXimation; QG: Quantile Graph; MTF: Markov Transition Field; BDNF: Brain-Derived Neurotrophic Factor; SHAM: Control condition without treatment.

To further assess discriminative capability, ROC curves were analyzed for each QTN method using gri-dpartitioned AIP time series (Figure 11). GAF maintained an AUC of 1.00, indicating flawless classification. QG and MTF also achieved near-perfect AUC values of 0.99, while SAX exhibited slightly lower but still robust discriminative power (AUC = 0.98). These ROC results corroborate the findings from the confusion matrix, reinforcing the superior performance of GAF and MTF representations under grid-partitioned AIP signals.

**FIG. 11:**
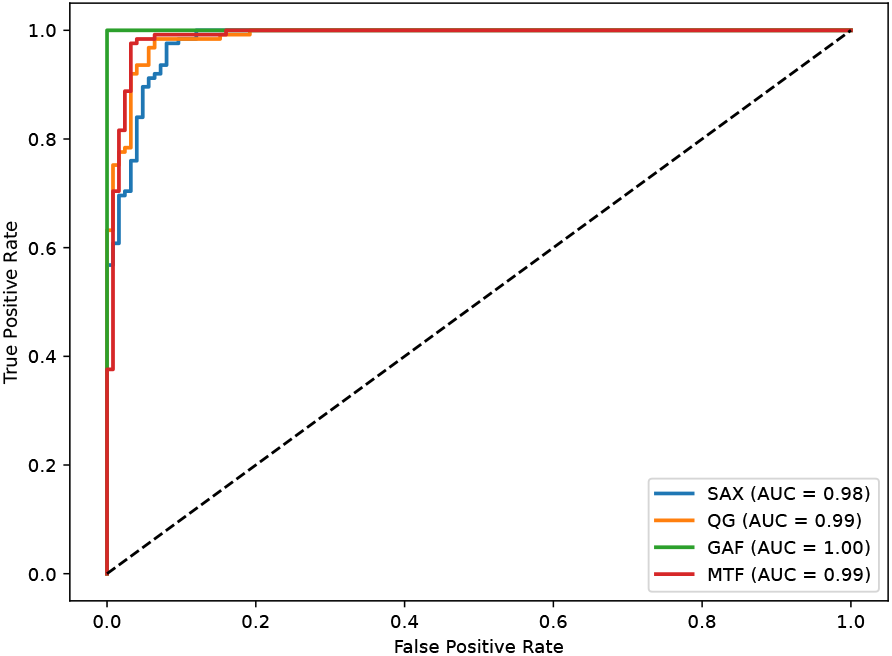
Grid-based partitioning ROC curves for each QTN method using 28 Hz. GAF achieved perfect classification (AUC = 1.00), while QG and MTF displayed near-perfect discrimination (AUC = 0.99). SAX maintained a high AUC of 0.98, demonstrating the overall robustness of grid-partitioned signal representations. QTN: Quantile-based Transition Network; AIP: Averaged Intensity Pixel; GAF: Gramian Angular Field; QG: Quantile Graph; MTF: Markov Transition Field; SAX: Symbolic Aggregate approXimation; AUC: Area Under the Curve.

Furthermore, the learning curves presented in Figure 12 illustrate the training and validation accuracies across epochs for each QTN method derived from grid-partitioned AIP time series. All methods exhibited stable convergence, with minimal discrepancies between training and validation accuracies, indicating effective generalization and negligible overfitting. Remarkably, the GAF representation achieved perfect classification accuracy in the early epochs, reflecting its capacity to capture highly distinctive temporal structures that facilitate rapid learning. SAX, QG, and MTF representations also demonstrated robust performance, converging to final validation accuracies of 0.928, 0.94, and 0.968, respectively, further validating the effectiveness of spatially localized time series descriptors in enhancing classification outcomes.

**FIG. 12:**
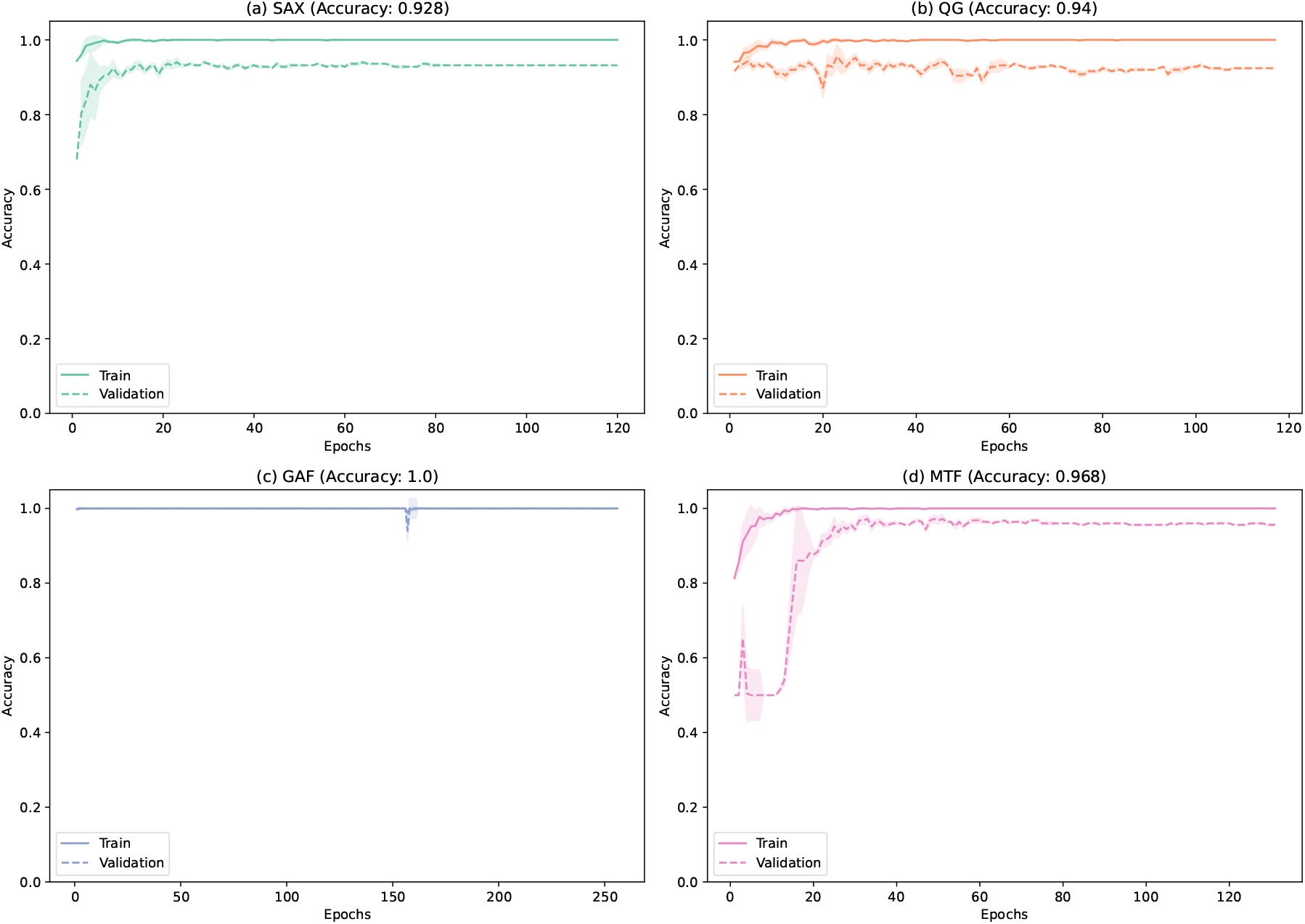
Grid-based partitioning learning curves for QTN methods using 28 Hz. Training (solid lines) and validation (dashed lines) accuracies are plotted across epochs for each method: (a) SAX, (b) QG, (c) GAF, and (d) MTF. Shaded regions represent the standard deviation, reflecting variability across training iterations. All methods achieved high final accuracies, with GAF reaching perfect classification. QTN: Quantile-based Transition Network; AIP: Averaged Intensity Pixel; SAX: Symbolic Aggregate approXimation; QG: Quantile Graph; GAF: Gramian Angular Field; MTF: Markov Transition Field.

To assess whether the CNN model’s high classification accuracy on Grid-AIP representations was robust to input perturbations, we evaluated performance degradation under progressive Gaussian noise addition similar to the previous subsection. As shown in Supplementary VII in Figures S1–(b), the model exhibited resilience, maintaining an AUC above 0.9 up to a noise standard deviation of approximately 2.0. Beyond this point, performance declined gradually as noise dominated the input signal. These results reinforce that the model’s discriminative capacity stems from meaningful activity patterns extracted through grid-based decomposition, rather than reliance on noise-driven artifacts.

### C. Validation of the model with the second experiments

As described in Section II, model generalization was assessed using recordings from a second, independent experiment performed on different biological samples prepared under identical conditions. This dataset, preprocessed with the grid-based strategy identified as most effective during training, was held out a priori and used exclusively to evaluate the external performance of the pre-trained CNN models. As summarized in Table II, the GAF representation achieved the highest performance, with an AUC of 0.99 and balanced accuracy, precision, and recall. Other QTN methods also performed strongly: SAX (AUC = 0.96), MTF (0.97), and QG (0.93). These results confirm the robustness of the QTN-based CNN framework, particularly GAF, in generalizing across datasets and capturing discriminative temporal dynamics beyond simple amplitude-based measures. To further illustrate classifier behavior, confusion matrices for each method are shown in Figure 13, demonstrating high true positive and true negative rates across QTN representations.

**FIG. 13:**
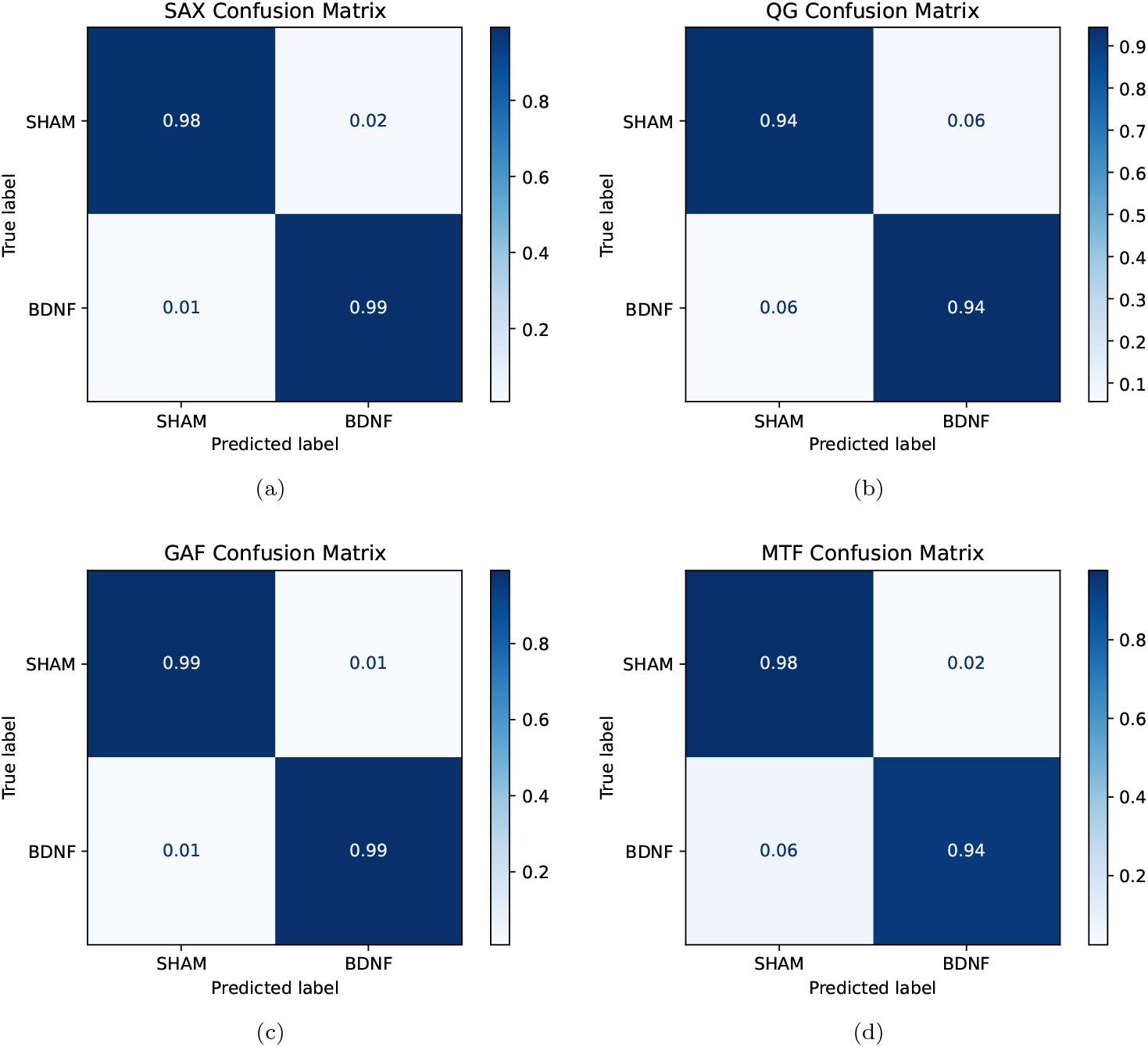
Confusion Matrices from Independent Validation using Grid-based partitioning and 28 Hz. Each panel shows the confusion matrix for a specific QTN-based CNN model evaluated on an independent dataset using grid-based preprocessing. GAF achieved the most balanced and highest classification accuracy, followed by SAX, MTF, and QG. Values are normalized. QTN: Quantile-based Time Series Representation; SAX: Symbolic Aggregate approXimation; QG: Quantile Graph; GAF: Gramian Angular Field; MTF: Markov Transition Field.

## VI. DISCUSSION

To comprehensively assess the influence of spatial resolution, acquisition frequency, and signal decomposition strategies on classification performance, we systematically compared two segmentation-agnostic preprocessing methodologies: Whole-Image Averaging (AIP) and Grid-Based Partitioning (Grid-AIP). For each strategy, we evaluated four Quantile-based Time-series (QTN) representations (SAX, QG, GAF, and MTF) using standard classification metrics (AUC, accuracy, precision, and recall). The corresponding summaries are reported for the whole-image analyses in Figures 4–6 and for the grid-based analyses in Figures 9–12. This design removes the dependence on single-cell segmentation and isolates the contributions of spatial granularity and sampling rate to CNN performance.

**TABLE II:** Independent validation performance using Grid-based partitioning and 28 Hz. All recordings were preprocessed using the grid-based partitioning strategy—the best-performing method identified in the training phase—and evaluated using CNN models trained on the original dataset. GAF yielded the highest performance, confirming its robustness across datasets. QTN: Quantile-based Transition Network; SAX: Symbolic Aggregate approXimation; QG: Quantile Graph; GAF: Gramian Angular Field; MTF: Markov Transition Field; AUC: Area Under the Receiver Operating Characteristic Curve.

| QTN Method | Accuracy | Precision | Recall | AUC |
| --- | --- | --- | --- | --- |
| SAX | 0.98 | 0.98 | 0.99 | 0.96 |
| QG | 0.94 | 0.93 | 0.94 | 0.93 |
| GAF | 0.99 | 0.99 | 0.99 | 0.99 |
| MTF | 0.95 | 0.97 | 0.93 | 0.97 |

Our analysis revealed that acquisition frequency played a critical role in classification outcomes. As illustrated in Figure 4 and Figure 5, a 28 Hz acquisition frequency consistently outperformed 4 Hz across all QTN methods, achieving near-perfect AUC values—particularly for GAF and QG representations. This suggests that higher temporal resolution facilitates the capture of subtle dynamic features associated with BDNF treatment, thereby improving discriminative performance.

Grid-AIP, in particular, achieved the highest classification scores, demonstrating that localized signal decomposition at an intermediate spatial scale (16×16 pixel grids) offers a more effective balance between granularity and signal robustness. This approach preserved critical spatial dynamics while mitigating segmentation artifacts, leading to superior model performance. Similarly, AIP maintained strong classification capabilities, underscoring the effectiveness of global intensity patterns, especially when fine segmentation is infeasible or unreliable due to image quality constraints.

Previous studies employing CNNs for calcium imaging classification have primarily relied on feeding raw pixel intensities from individual imaging frames directly into the network, as discussed in Section I. While effective at capturing spatial information, this approach is computationally intensive and does not scale well with large datasets. In contrast, our framework leverages QTN representation applied to both AIP and Grid-AIP, offering a more scalable and efficient alternative. Notably, all QTN representations used here employ a discretization parameter, as established in the work of [24], which determines the number of intervals into which each time series is divided.

As noted in the Introduction I, CNN-based analyses—especially with high-resolution or volumetric inputs—incur substantial computational and memory costs. A practical advantage of the QTN approach is controlled dimensionality reduction. Rather than feeding entire fluorescence sequences of length *T* into the model, each trace is summarized as a *Q* × *Q* matrix with *Q ≪T* (see Section IV A). In our dataset, recordings with 3,356 frames are mapped to 30 × 30 (900 values), and those with 460 frames to 15 × 15 (225 values). This fixed-size representation reduces memory and computation, simplifies batching across recordings of different lengths, and can lower overfitting risk by constraining the hypothesis space—while preserving ordinal/transition structure that is critical for neuronal dynamics. The stability we observe under noise perturbations and across sampling rates indicates that this compression retains task-relevant temporal information.

Among QTN methods, the GAF consistently demonstrated the most stable and robust classification performance across all preprocessing strategies, achieving AUC values exceeding 0.9. As evidenced by the learning curves shown in Figures 8 and 12, CNN models trained on GAF representations converged to optimal performance within only a few epochs. This rapid convergence suggests that GAF encodes highly discriminative temporal features, enabling the network to efficiently learn class boundaries. While this behavior likely reflects GAF’s capacity to project meaningful temporal patterns into a separable feature space, it may also partially result from the relatively low intra-class variability or overall simplicity of the current classification task. To test the generalizability of this approach, future work will evaluate its performance across more complex datasets, including recordings from distinct tissue types and experimental conditions with higher biological heterogeneity. Importantly, the robustness analysis under progressive Gaussian noise perturbations (Figure S1-(a) and (b)) confirms that the CNN’s discriminative capacity is not driven by overfitting or memorization but instead relies on temporally coherent and biologically relevant signal structure.

Furthermore, to further confirm the generalization capability of the proposed framework, we evaluated the CNN models on an independent dataset using the optimal grid-based preprocessing strategy. As detailed in Table II and visualized in Figure 13, all QTN representations maintained high classification performance, with GAF achieving an outstanding AUC of 0.99. SAX and MTF also demonstrated strong generalization with AUCs of 0.96 and 0.97, respectively, while QG maintained solid performance at 0.93. The confusion matrices highlight consistent classification accuracy across both classes, underscoring the model’s reliability beyond the training distribution. These findings reinforce the robustness and practical applicability of the QTN-based CNN approach in real-world scenarios and further validate the superiority of GAF in capturing class-distinct temporal structures. A key advantage of QTN lies in its ability to encode temporal information from a single fluorescence time series into a matrix suitable for CNN architectures, effectively preserving temporal ordering and local signal structures while simplifying the preprocessing pipeline. To our knowledge, this is the first study to apply QTN-based CNN classification to calcium imaging datasets, specifically for distinguishing between BDNF-treated and SHAM samples. This provides a scalable and accurate framework for future neuropharmacological analyses. Nonetheless, despite the strong classification performance, a key limitation of this approach lies in the challenge of biological interpretability. The internal representations learned by the CNN—and, by extension, the structure of the QTN matrices—remain largely opaque in terms of their correspondence to underlying neurophysiological mechanisms. To address this, future work will focus on incorporating graph-theoretical descriptors (e.g., node degree [59], clustering coefficient [60], modularity [59]) extracted from QTN-derived networks and investigating how changes in network topology relate to biologically meaningful alterations in cellular dynamics.

Table III provides a consolidated overview of the pre-processing approaches, highlighting the trade-offs between spatial resolution, classification performance (as obtained in the current results), and computational cost. Grid-AIP emerged as the optimal trade-off strategy—combining high classification accuracy with manageable computational demands—while Whole-Image Averaging (AIP) offered the lowest computational cost but at the expense of spatial detail. The comparative analysis demonstrates that increasing spatial granularity through grid partitioning enhances the discriminative power of QTN-based representations without imposing excessive processing requirements. This suggests that a moderate spatial resolution, as achieved with the grid approach, is sufficient to capture meaningful fluorescence dynamics, preserving both computational efficiency and interpretability.

## VII. CONCLUSIONS AND FUTURE WORK

This study presents a novel, scalable, and computationally efficient framework for classifying calcium imaging dynamics. It couples QTN representations with convolutional neural networks and is systematically evaluated across multiple spatial resolutions and acquisition frequencies. By compressing each fluorescence time series of length *T* into a fixed *Q × Q* matrix (*Q ≪ T* ), the method reduces input dimensionality, while standardizing inputs across recordings. Our findings demonstrate that global and grid-based preprocessing strategies—particularly AIP and Grid-AIP—consistently achieved high classification performance with minimal computational overhead. Among all tested QTN representations, the GAF emerged as the most robust and stable descriptor, effectively capturing treatment-induced neuronal dynamics across varying spatial and temporal conditions. Notably, classification performance remained high even at lower acquisition frequencies, underscoring the framework’s resilience to reduced temporal resolution.

To our knowledge, this is the first application of quantile-based network representations in combination with CNNs for the classification of calcium imaging data. The ability of this framework to accurately distinguish between treated and control samples using only a single time series per region highlights its potential as a generalizable and efficient tool for functional imaging analysis. Future work will focus on expanding this methodology to include additional neuroactive compounds with diverse mechanisms of action, such as psychedelics, neuromodulators, and neurotoxins, thereby enabling broader applications in drug screening, disease modeling, and systems neuroscience. In parallel, we plan to adapt the framework to more challenging segmentation scenarios, including those with low contrast and densely packed tissues. Lastly, we will use the new, longer version of the QG method that was just made to change grayscale images into a structured network format [61]. These approaches will help us analyze and classify the dynamics of calcium imaging, giving us more information about neuronal activity patterns.

## NOMENCLATURE

**Abbreviations**

AIP: Averaged Intensity Pixel
AUC: Area Under the ROC Curve
BDNF: Brain-Derived Neurotrophic Factor
CNN: Convolutional Neural Network
CV: Coefficient of Variation
DC: Direct-current component
DAIP: First temporal derivative of AIP
FOV: Field of View
GAF: Gramian Angular Field
Hz: Hertz (cycles per second)
IQR: Interquartile Range
LF: Low-Frequency band
MTF: Markov Transition Field
QG: Quantile Graph
QTN: Quantile-Based Time-Series Network
rFFT: real Fast Fourier Transform
ROC: Receiver Operating Characteristic
ROI: Region of Interest
SAX: Symbolic Aggregate approXimation
SDAIP: Second temporal derivative of AIP
SHAM: Control condition without treatment

**Symbols**

*Q*: Number of quantile bins
*T*: Length (number of frames) of the time series
*f*_*s*_: Sampling rate (Hz)
Δ*f*: Frequency resolution, *f*_*s*_*/T*
*F* (*k*): Complex Fourier coefficient at bin *k*
*P*_LF_: Low-frequency spectral power

## ACKNOWLEDGEMENTS

CT gratefully acknowledges financial support from the Deutsche Forschungsgemeinschaft (DFG, German Research Foundation)—project number 498176770 (light sheet microscope project); the Deutsche Forschungsgemeinschaft (DFG) in the frame of “Untersuchung der Wirkung von psychedelischen Substanzen auf neuronale dreidimensionale Zellkulturen” (grant TH 1448/5-1), the Bundesministerium für Forschund, Technologie und Raumfahrt (BMFTR) in the frame of iNeuTox (grant 13FH516IX6), and the Zentrum für Wissenschaftliche Services und Transfer (ZeWiS), Germany. ASLOC acknowledges support from the Fundação de Amparo à Pesquisa do Estado de São Paulo (FAPESP), grant 2023/06563-9. Furthermore, the authors acknowledge support from the BMBF project ESTRANGE (grant 02NUK081B).

## DECLARATION OF GENERATIVE AI AND AI-ASSISTED TECHNOLOGIES IN THE WRITING PROCESS

During the preparation of this work the authors used ChatGPT in order to improve the readability of this paper. After using this tool, the authors reviewed and edited the content as needed and take full responsibility for the content of the published article.

## DATA AVAILABILITY

Data requests can be directed to the corresponding authors and will be provided upon reasonable request, subject to institutional and ethical guidelines.

## AUTHOR CONTRIBUTIONS

Conceptualization: CLA, CT. Data curation: SH, HD. Formal analysis: CLA. Funding acquisition: CT. Investigation: CLA, LS, SH, MM, ASLOC, FAR, CT. Methodology: CLA. Project administration: CT. Resources: CT. Software: CLA. Supervision: CT, FAR. Validation: CLA, LFS, SH, MM, ASLOC, FAR, CT. Visualization: CLA, ASLOC. Writing – original draft: CLA, SH, MM, CT. Writing – review and editing: CLA, LFS, SH, MM, ASLOC, FAR, CT.

**TABLE III:** Comparison of Preprocessing Strategies. Overview of spatial resolution, classification performance (as observed in the current results), and computational cost for Whole-Image Averaging (AIP) and Grid-Based Partitioning (Grid-AIP). Performance reflects the classification accuracy and AUC obtained in this study, while cost refers to the relative computational resources required for preprocessing and feature extraction. AIP: Averaged Intensity Pixel; Grid-AIP: Grid-Based Averaged Intensity Pixel; AUC: Area Under the Curve.

| Preprocessing Approach | Spatial Resolution | Performance | Computational Cost |
| --- | --- | --- | --- |
| AIP | Global | High | Low |
| Grid-AIP | Medium (16×16 grids) | Highest | Moderate |

## S1. ROBUSTNESS OF CNN

To examine whether the convolutional neural network (CNN) maintained its discriminative ability under signal perturbations, we performed a robustness analysis by adding Gaussian noise of increasing standard deviation to the QTN-based input matrices. For both the Whole-Image Averaging and Grid-Based Partitioning approaches, we quantified the degradation in classification accuracy using the area under the ROC curve (AUC) as a function of noise intensity as shown in Figure S1. This analysis assesses whether the model relies on meaningful temporal–spatial patterns rather than noise-driven artifacts.

**FIG. S1:**
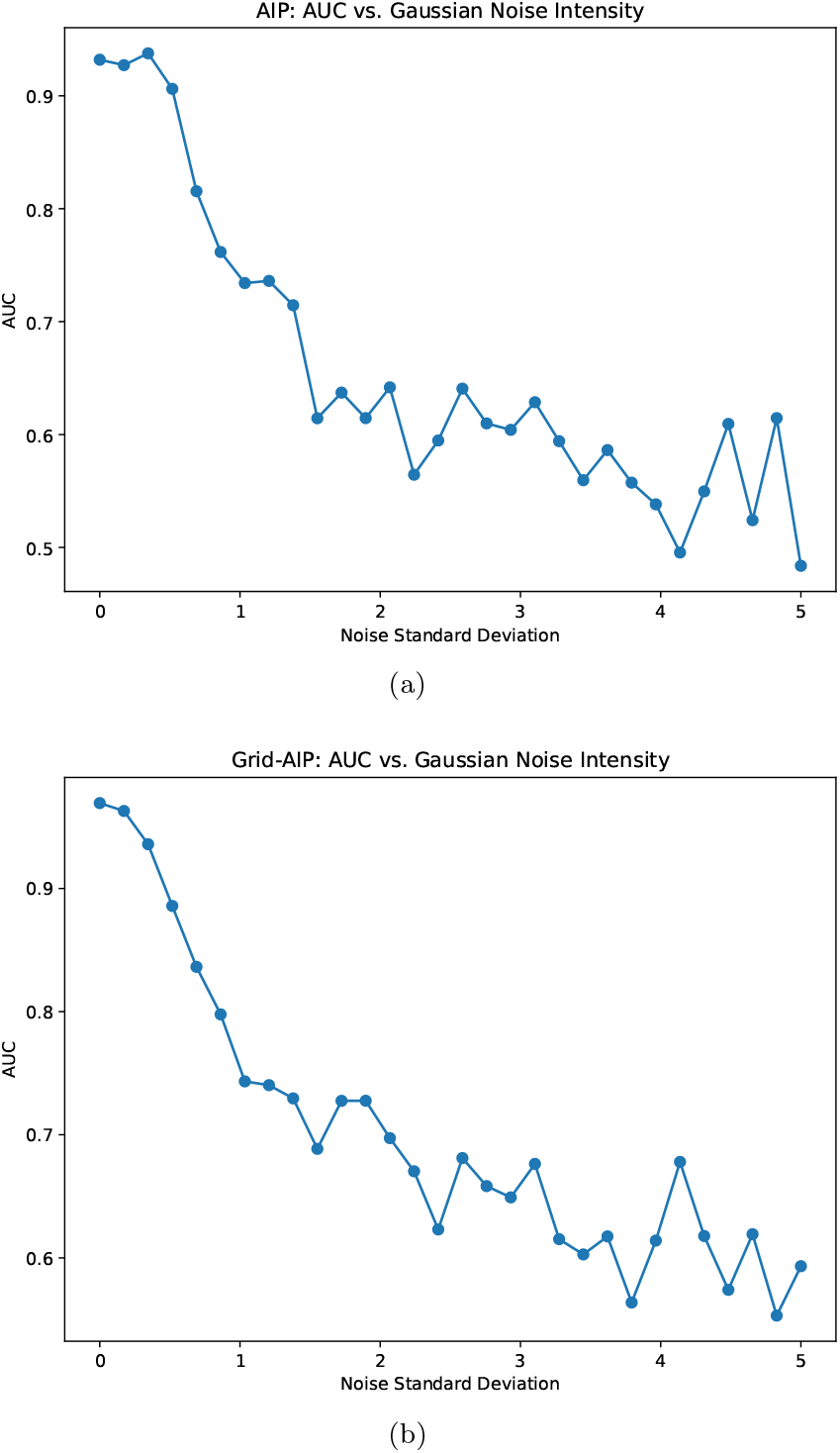
Robustness of GAF representations and CNN to Gaussian noise. Both panels show the degradation of classification performance (AUC) as a function of increasing Gaussian noise (standard deviation) added to the QTN matrices derived from Averaged Intensity of Pixel (AIP) signals. The CNN remained resilient up to moderate noise levels, with a gradual decline in performance as noise dominated the input. AUC: Area Under the Curve; GAF: Gramian Angular Field; QTN: Quantile-based Time-series representation; AIP: Averaged Intensity of Pixel.

